# FADD- and RIPK3-mediated cell death ensures timely clearance of wound macrophages and promotes wound healing

**DOI:** 10.1101/2023.03.29.534669

**Authors:** Louise Injarabian, Sebastian Willenborg, Daniela Welcker, Manolis Pasparakis, Hamid Kashkar, Sabine A Eming

## Abstract

Cells of the monocyte/macrophage lineage are an integral component of the body’s innate ability to restore tissue function after injury. In parallel to mounting an inflammatory response, clearance of monocytes/macrophages from the wound site is critical to re-establish tissue functionality and integrity during the course of healing. The role of regulated cell death in macrophage clearance from damaged tissue and its implications for the outcome of the healing response is little understood. Here, we explored the role of macrophage-specific FADD-mediated cell death on *Ripk3^-/-^* background in a mechanical skin injury model in mice. We found that combined inhibition of RIPK3-mediated necroptosis and FADD-caspase-8-mediated apoptosis in macrophages profoundly delayed wound healing. Importantly, RIPK3 deficiency alone did not considerably alter the wound healing process and macrophage population dynamics, arguing that inhibition of FADD-caspase-8-dependent death of macrophages is primarily responsible for the delayed wound closure. Notably, TNF blockade reversed the accumulation of Ly6C^high^ macrophages induced by combined deficiency of FADD and RIPK3, indicating a critical dual role of TNF-mediated pro-survival and cell death signalling, particularly in this highly pro-inflammatory macrophage subset. Our findings reveal a previously uncharacterized cross-talk of inflammatory and cell death signalling in macrophages in regulating repair processes in the skin.

## Introduction

The skin provides a life-sustaining structural and immunological outer barrier of the body, which requires continuous repair and regeneration. Skin injury extending into the dermis induces a complex, dynamic cellular program proceeding in sequential stages of tissue growth and differentiation (Gurtner, 2008). The healing response in skin due to excision injury is initiated by local inflammation, followed by the formation of vascularized granulation tissue characterized by fibroblast-myofibroblast differentiation, resolution of inflammation and activation and deposition of provisional extracellular matrix (ECM) that ultimately converts into a collagenous fibrous scar tissue. Repair of the dermal tissue is paralleled by regeneration of the overlying epidermis leading to wound closure.

Cells of the monocyte-macrophage lineage are an integral component of an effective repair response; their inappropriate activation has emerged as one of the pivotal mechanisms underlying the pathophysiology of repair (Murray, 2011; Eming, 2010; Eming, 2014; Ding, 2015). Blood monocytes that are recruited to the site of tissue damage sense a variety of environmental cues of injured tissue and integrate those into a host protective wound healing response (Willenborg, 2012; Knipper, 2015; Kiesewetter, 2019; Hadrian, 2023). In addition to the requirement to sense and respond rapidly to distinct damage signals, macrophages need to be cleared from the wound site at later time points in order to govern the return to tissue homeostasis (Eming, 2014; Knuever, 2015; Eming, 2017; Do, 2018). The dominant fate of injury-associated monocytes/macrophages at the wound site during resolution of inflammation is unclear and under debate (Majka, 2010; Sinha, 2018, Li, 2021). Recent evidence, particularly in tissues other than skin, suggests a critical role of different forms of myeloid cell death in tissue homeostasis and repair (Moriwaki, 2014; Bleriot, 2015). How different modes of macrophage death contribute to resolution of inflammation and the repair response is little understood.

By transcriptional profiling of early and late-stage wound macrophages we recently uncovered that early stage/type-1 towards late stage/type-2 macrophage activation dynamics are paralleled by TNF-related activation profiles (Willenborg, 2021; Sanin, 2022). Particularly early stage wound macrophages revealed a TNF-mediated cell death associated gene profile including increased expression of *Tnf, Ripk1, Fadd, Ripk3, Mlkl* (Willenborg, 2021; Sanin, 2022). Based on these findings, we hypothesized an important role for regulated extrinsic cell death in wound macrophages contributing to resolution of inflammation and repair. To examine the role of apoptosis and necroptosis in myeloid cells during the course of skin wound healing we subjected wild-type, global RIPK3^-/-^ and FADD^MKO^RIPK3^-/-^ mice (myeloid cell-restricted *Fadd* deletion on *Ripk3^-/-^* background) to excision skin injury. Our findings highlight FADD/RIPK3-dependent cell death as novel mechanism required for timely clearance of a pro-inflammatory macrophage subset from damaged skin. In addition, our findings reveal an important role of TNF in regulating this process. Together, our findings contribute to a deeper mechanistic understanding of anti-TNF-pro repair activities in tissue destructive inflammatory conditions that might ultimately guide the development of therapeutic approaches to ameliorate wound healing disorders in patients.

## Materials and methods

### Animals

Mouse strains used in this study are listed in **Table 1**. Animal housing and all the experimental procedures were authorized by the North Rhine–Westphalian State Agency for Nature, Environment, and Consumer Protection (LANUV, North Rhine– Westphalia, Germany) and the University of Cologne (81-02.04.2020.A466). All mouse lines used did not suffer from their gene modifications. Only mice which appeared healthy and were not involved in previous procedures were used for experiments. Mice were housed 2-5 per cage in a temperature (22-24°C) and humidity (relative humidity of 45-65 rH) controlled colony room, maintained on a 12 h light/dark cycle, with standard food (Altromin) and water provided *ad libitum*. The hygiene monitoring took place quarterly according to the recommendations of the FELASA (Federation of European Laboratory Animal Science Associations) for the health monitoring of laboratory animal facilities. The hygiene status of the animal husbandry was SPF.

**Table 1.** Mouse strains.

| **Experimental models (strain)** | **Reference** | **Link** |
| --- | --- | --- |
| C57BL/6J mice | Charles River | https://www.criver.com/products-services/find-model/jax-c57bl6j-mice?region=23 |
| *Fadd<sup>fl/fl</sup>Ripk3<sup>-/-</sup>* mice | Newton, 2004;<br>Mc Guire, 2010 | N/A |
| *Cx3cr1*-cre mice | Yona, 2013 | N/A |
| *Fadd<sup>fl/fl</sup>Ripk3<sup>-/-</sup> Cx3cr1<sup>Cre</sup>* mice | (generated in this study) | N/A |

*FADD^MKO^RIPK3^-/-^* mice were generated by inter-crossing *Fadd^fl/fl^* (Mc Guire, 2010), *Ripk3^-/-^* mice (Newton, 2004) and *Cx3cr1^Cre^* mice (Yona, 2013) (C57BL/6N background). All mice used in the experiments were generated by in-house mating. The sequences of the primers used to genotype the mice and to verify Cre-mediated recombination are provided in **Table 2**. For wound healing studies, 8-to-14-week-old male or female mice were used for experiments. Animals were first allocated to different groups based on their genotype and then randomized for different time points of analysis. For TNF signaling inhibition experiments, mice were treated with Etanercept (i.p., 12 µg/g body weight in 200µL NaCl 0.9%) in a 3-day interval starting 3 days prior to wounding. After wounding the animals were single housed.

**Table 2.**
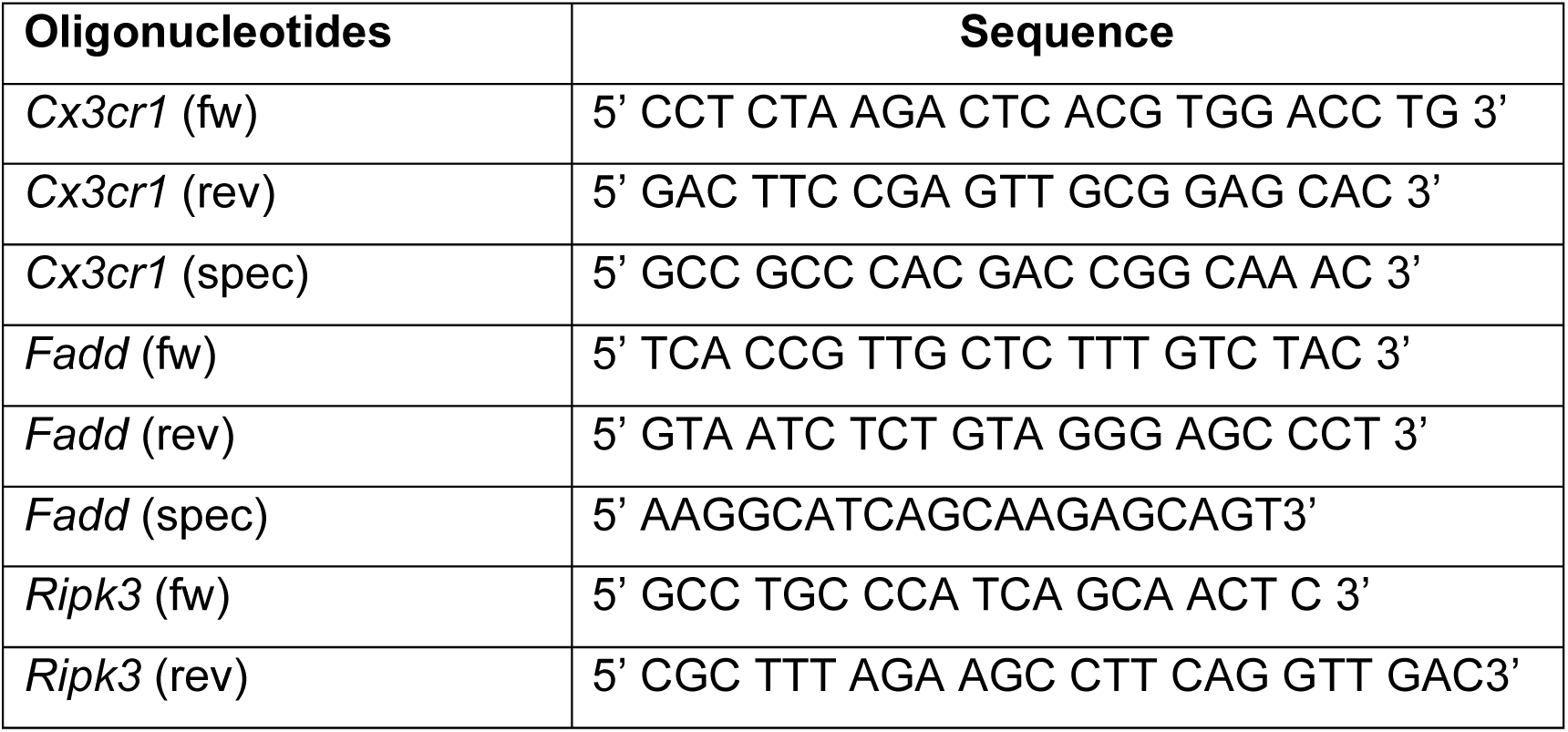
Primer sequences for genotyping.

### Excisional punch injury

Mice were anesthetized by intraperitoneal injection of 100 mg/kg body weight Ketavet (Pfizer) and 10 mg/kg body weight Rompun 2% (Bayer). The back skin was shaved using an electric shaver and disinfected with 70% ethanol. Full-thickness punch biopsies were created on the back using a standard biopsy puncher (Stiefel). For histological analysis, wounds were excised at different time points after injury and bisected in the caudocranial direction and processed following an established protocol (Lucas, 2010; Willenborg, 2012; Knipper, 2015). The tissue was either fixed overnight in Roti®Histofix or embedded in O.C.T. compound (Fisher Scientific) and stored at - 80°C.

### Morphometric analysis

Morphometric analysis was performed on H&E-stained paraffin tissue sections using light microscopy equipped with a KY-F75U digital camera (JVC) at various magnifications (Leica DM4000B, Leica Microsystems; Diskus 4.50 software) as described previously (Lucas, 2010; Willenborg, 2012; Knipper, 2015). The extent of granulation tissue formation was assessed. The gap between the epithelial tips was determined as a measure of wound closure. Analysis was performed in a blinded manner.

### Immunostaining

For ɑSMA immunofluorescence staining, cryostat sections (10μm thick) from O.C.T. embedded tissues were fixed in acetone (stored at -20°C) for 5 min, washed in PBS 3 times, and blocked in 10% normal goat serum in PBS for 1h at RT. Slides were incubated with anti-ɑSMA-Cy3 (Sigma Aldrich, C6198), diluted 1:250, and DAPI, 1µg/mL for 2h at RT in antibody diluent (DCS Innovative Diagnostik-Systeme). After three washes, slides were covered with coverslips in mounting medium (Prolong^TM^ Gold Antifade Mountant, Thermo Fisher, P36930). Stained sections were analyzed using the Keyence BZ-9000 fluorescence microscope and the BZ-II viewer software and analyzed with ImageJ.

### Flow cytometry & FACS sorting

Single-cell suspensions of wound tissue were prepared by a combination of enzymatic digestion (Liberase Blendzyme, Roche Applied Science) and mechanical disruption (Medimachine System, BD Biosciences) as described previously (Willenborg, 2021). Briefly, excised wound tissue was sectioned with a scalpel, placed in DMEM medium supplemented with 30mg/mL Liberase TM Research Grade (Roche), and incubated at 37°C for 90min (shaking). Digested wound tissue was mechanically disrupted for 5 min using the Medimachine System (BD Biosciences). Cells were passed through 70µm and 40µm cell strainer and washed with ice-cold PBS/1% BSA/2mM EDTA. Leukocytes were isolated from whole blood by hypotonic lysis of erythrocytes using Red Blood Cell Lysis Solution (Miltenyi Biotec) followed by ice cold PBS/1% BSA/2mM EDTA washes. Fc receptors were blocked with anti-CD16/CD32 (diluted 1:50, Thermo Fisher Scientific) and cells were stained with cell surface antigens, namely fluorescein isothiocyanate (FITC)- or Pacific blue-conjugated anti-CD45 (diluted 1:200, Thermo Fisher Scientific), allophycocyanin (APC)-conjugated or APC-Cy7-conjugated anti- CD11b (diluted 1:300, Thermo Fisher Scientific), phycoerythrin (PE)-conjugated anti- F4/80 (diluted 1:50, Bio-Rad Laboratories), and PE-Cy7-conjugated Ly6C (diluted 1:400, BD Biosciences) and APC-Cy7-conjugated Ly6G (diluted 1:400, BD Biosciences). To assess apoptosis, cells were first stained with surface markers and then stained with Pacific blue-conjugated Annexin V (Thermo Fischer Scientific, R37177) for 25 min at RT in Annexin V binding buffer (Thermo Fischer Scientific, V13246). Dead cells were excluded using 7-AAD (7-Aminoactinomycin D, Thermo Fisher Scientific, 00-6993-50). Cells were analyzed using a FACSCanto II flow cytometer (BD) and sorted using a FACSAria III cell sorting system (BD Biosciences) equipped with FACSDiva Version 6.1.1 software (BD Biosciences). FACS data were further analyzed by Flowjo Version 10.8.2 (FlowJo).

### Quantitative real-time PCR

For wound monocytes/macrophages and blood monocytes, cytometry-sorted 7-AAD^-^ CD45^+^CD11b^+^F4/80^+^ cells were lysed in 350µL RLT lysis buffer (Qiagen, 79216). At 4 days post injury, 4 wounds were pooled, and at 14 days post injury, 6-8 wounds were pooled in order to obtain 30-50 000 monocytes/macrophages for analysis. Total RNA was isolated using the RNeasy Plus Micro or Mini Kit (Qiagen; 74104 /74034) according to the manufacturer’s instructions. Reverse transcription of isolated RNA was performed using the High-Capacity cDNA RT Kit (Thermo Fisher Scientific, 4368814). Amplification reactions (triplicates) were set up using the PowerSYBR Green PCR Master Mix (Thermo Fisher Scientific, 4367659) and qRT-PCR was performed using the QuantStudio™ 5 Real-time PCR system (Thermo Fisher Scientific). The comparative method of relative quantification (2^-ΔΔCt^) was used to calculate the expression level of the target gene normalized to a housekeeping gene *(Rps29*). All primer sequences are listed in **Table 3**.

**Table 3.**
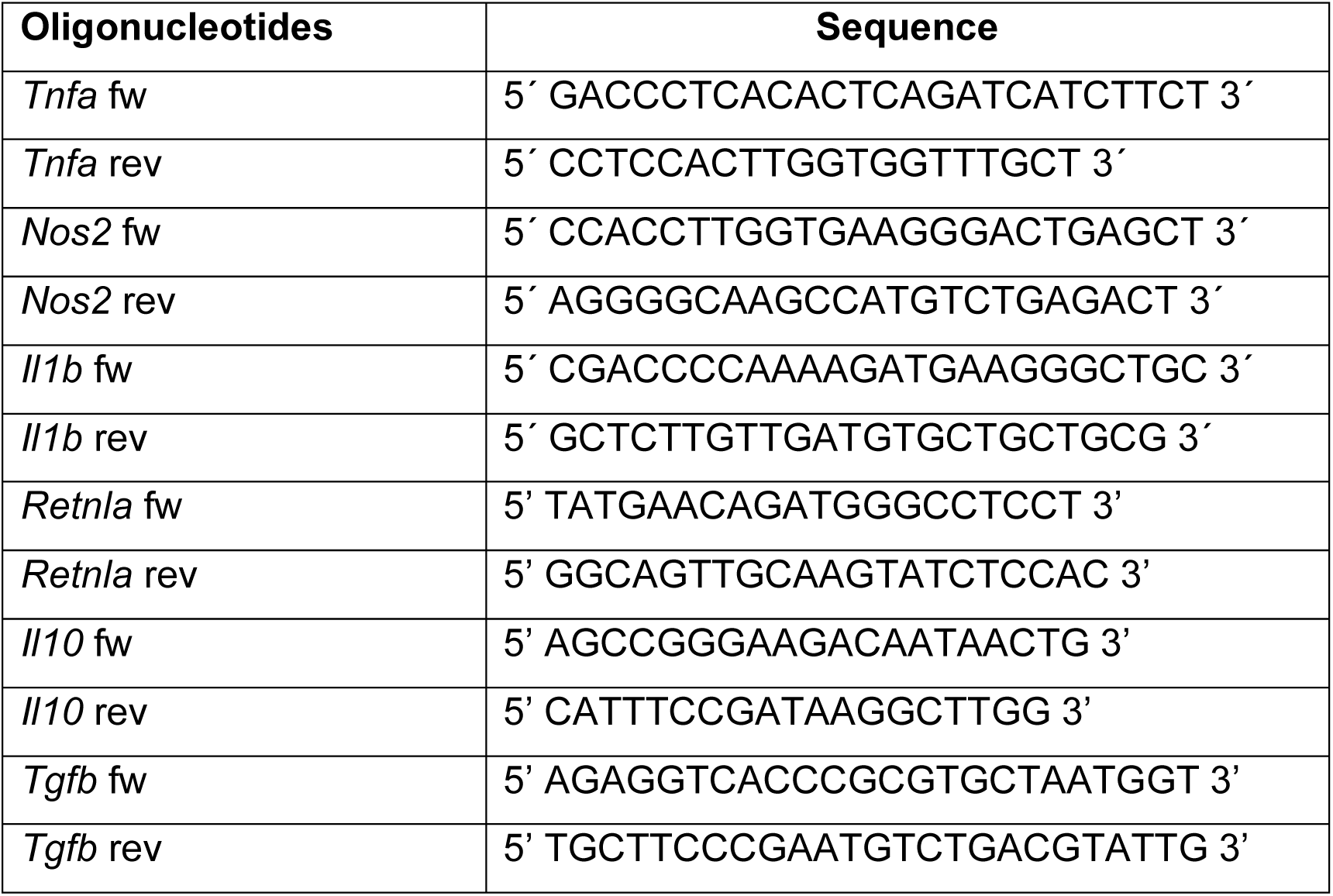
Primer sequences for qRT-PCR.

### Statistics

Parametric methods, i.e., Student’s unpaired two-tailed t- test (when 2 groups were compared) or one-way ANOVA with Tukey’s post-test (when more than 2 groups were compared) were used (GraphPad Prism 9 software). In case of not assumed equal variances, a Welch correction was performed. All *P value*s are reported in the figure legends. Results are considered significant when *P* < 0.05.

## Results

### Cell death regulates the number of monocytes/macrophages in physiological skin repair

To analyze the dynamics of the cellular composition of immune and non-immune cells over the time course of physiological skin repair, we inflicted full-thickness excisional skin wounds on the back skin of C57BL/6N wild-type mice and subjected wound cell suspensions in the early inflammatory phase (4 days post-injury [dpi]), the mid tissue forming phase (7 dpi) and the late scar forming phase (14 dpi), to flow cytometry. Analysis revealed a high proportion of CD45^+^ leukocytes at 4 dpi (87.9% ± 9.9%), which gradually declined towards the late phase of healing (7 dpi: 73.6% ± 11.8%, 14 dpi: 57.5% ± 6.6%); in parallel, consistent with a productive repair process of the damaged tissue, the fraction of CD45^-^ stromal cells (containing mainly keratinocytes, fibroblasts/myofibroblasts, endothelial cells) increased gradually **(Fig. 1A).**

**Figure 1.**
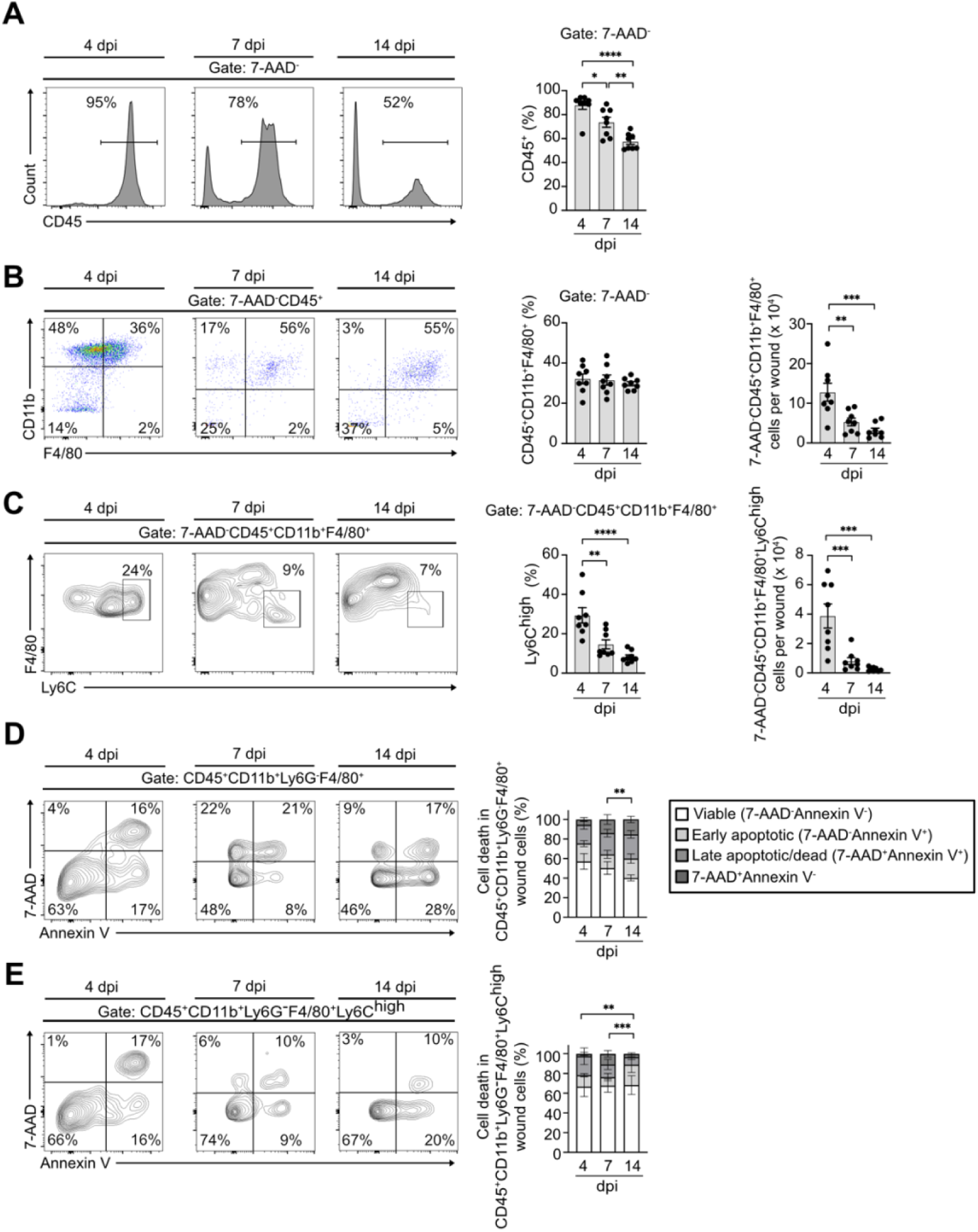
Cell death regulates monocyte/macrophage numbers in physiological skin repair. **A.** Wound cells isolated from wild-type mice at indicated days post injury (dpi) and analyzed by flow cytometry for CD45 expression. **B.** 7-AAD^-^CD45^+^ cells were gated and analyzed for CD11b and F4/80 expression. **C.** 7-AAD^-^CD45^+^CD11b^+^ F4/80^+^cells were gated and analyzed for Ly6C expression. (A-C) mean value +/- SEM is represented. Analysis of cell death by 7-AAD/Annexin V double staining in wound cells isolated from wild-type mice at indicated dpi (flow cytometry) in **D.** wound macrophages (CD45^+^CD11b^+^Ly6G^-^F4/80^+^) and in **E**. Ly6C^high^ inflammatory wound macrophages (CD45^+^CD11b^+^Ly6G^-^F4/80^+^Ly6C^high^). (D, E) mean value +/- SD is represented. *p<0.05, **p<0.01, ***p<0.001, ****p<0.0001.

We then focused our analysis on wound monocyte/macrophage population dynamics (wound monocytes/macrophages hereinafter defined as CD45^+^CD11b^+^F4/80^+^ cells). The relative number of CD45^+^CD11b^+^F4/80^+^ cells showed minor changes over the course of the healing response (4 dpi: 32.6% ± 6.9%, 7 dpi: 31.5% ± 7.1%, 14 dpi: 29.7% ± 2.9%) (**Fig. 1B**). As expected, the absolute number of viable wound monocytes/macrophages per wound significantly decreased over the course of repair (4 dpi: 128000 ± 64000; 7 dpi: 53000 ± 27000, 14 dpi: 31000 ± 18000) (**Fig. 1B**).

Within the wound monocyte/macrophage compartment we identified Ly6C^high^ cells, known to boost inflammation and angiogenesis, both processes essential for efficient tissue formation and timely healing (Willenborg, 2012). Confirming our previous findings (Willenborg, 2012), relative and absolute numbers of Ly6C^high^ monocytes/macrophages were highest in the inflammatory phase and gradually decreased in later phases of healing (relative: 4 dpi: 29.4% ± 11.0%, 7 dpi: 14.6% ± 6.5%, 14 dpi: 8.2% ± 3.0%; absolute: 4 dpi: 38000 ± 23000; 7 dpi: 8000 ± 7000, 14 dpi: 2000 ± 1000 (**Fig. 1C**). We asked whether cell death might contribute to the prominent decline of Ly6C^high^ wound monocytes/macrophages during healing progression. To assess cell death in wound monocytes/macrophages, we subjected wound single cell suspensions isolated from wild-type mice at different time points after injury to 7-AAD and Annexin V double staining, followed by flow cytometric analysis. As apoptotic polymorphonuclear cells (PMNs) give a strongly positive signal for Annexin V, we excluded PMNs from our analysis by gating CD45^+^CD11b^+^Ly6G^-^F4/80^+^ cells (**Fig. S1**).

Our analysis revealed the presence of a significant proportion of early apoptotic (7- AAD^-^Annexin V^+^) and late apoptotic (7-AAD^+^Annexin V^+^) CD45^+^CD11b^+^Ly6G^-^F4/80^+^ cells over the course of healing (**Fig. 1D**). Notably, the proportion of early apoptotic CD45^+^CD11b^+^Ly6G^-^F4/80^+^ wound cells increased significantly from 7 dpi to 14 dpi (early apoptotic at 7 dpi: 14.0% ± 4.1%, at 14 dpi: 19.8% ± 4.8%) (**Fig. 1D**). To investigate whether cell death occurs in the inflammatory Ly6C^high^ fraction of wound monocytes/macrophages, we gated CD45^+^CD11b^+^Ly6G^-^F4/80^+^Ly6C^high^ cells and analyzed 7-AAD and Annexin V fluorescence. Early and late apoptotic CD45^+^CD11b^+^Ly6G^-^F4/80^+^Ly6C^high^ cells were detected at all stages of repair (**Fig. 1E**). Notably, the proportion of early apoptotic CD45^+^CD11b^+^Ly6G^-^F4/80^+^Ly6C^high^ wound monocytes/macrophages was significantly higher at 14 dpi compared to 4 and 7 dpi (early apoptotic at 4 dpi: 11.6% ± 1.6%, at 7 dpi: 8.6% ± 3.7%, at 14 dpi: 20.7% ± 8.0%). Taken together, these findings suggest that apoptotic cell death plays a critical role in regulating relative and absolute numbers of pro-inflammatory Ly6C^high^ monocytes/macrophages in physiological repair.

### Modulating FADD-RIPK3 signaling in monocytes/macrophages impairs tissue repair

To elucidate the role of apoptosis and necroptosis in monocytes/macrophages during wound healing, we generated mice lacking FADD specifically in the monocyte/macrophage lineage in a *Ripk3*^-/-^ genetic background (*Fadd^fl/fl^Ripk3^-/-^ Cx3cr1^Cre/wt^*, hereafter abbreviated as FADD^MKO^RIPK3^-/-^) (**Fig. S2A**). The adaptor protein FADD (<u>F</u>as-<u>A</u>ssociated protein with <u>D</u>eath <u>D</u>omain) plays a central role in cell death ligand induced caspase-8 dependent apoptosis, whereas RIPK3 (<u>R</u>eceptor Interacting Serine/Threonine <u>K</u>inase <u>3</u>) is a key mediator of necroptosis (Pasparakis, 2015). Therefore, in FADD^MKO^RIPK3^-/-^ both extrinsic apoptosis and necroptosis are inhibited in monocytes/macrophages. The offspring were obtained in accordance with Mendelian laws (**Fig. S2B**) and developed normally, with no major spontaneous phenotypes in specific-pathogen-free (SPF) conditions. No significant premature death was noted. We confirmed an approximately 55% *Cx3cr1-*cre-mediated *Fadd* gene deletion and absence of *Ripk3* in wound macrophages of FADD^MKO^RIPK3^-/-^ mice (**Fig. 2A; Fig. S2C**). Because wound monocytes/macrophages are primarily derived from recruited blood monocytes (Willenborg, 2012), we evaluated the composition of the myeloid cell compartment in the blood of FADD^MKO^RIPK3^-/-^ mice compared to RIPK3^-/-^ mice using flow cytometry. No major differences between the lymphocyte (CD45^+^CD11b^-^SSC^low^), PMN (CD45^+^CD11b^+^Ly6G^+^SSC^high^) and Ly6C^high^ monocyte (CD45^+^CD11b^+^SSC^low^Ly6C^high^) compartments in the blood of naïve (not wounded) FADD^MKO^RIPK3^-/-^ mice compared to the RIPK3^-/-^ mice was observed (**Fig. S2D, E**).

**Figure 2.**
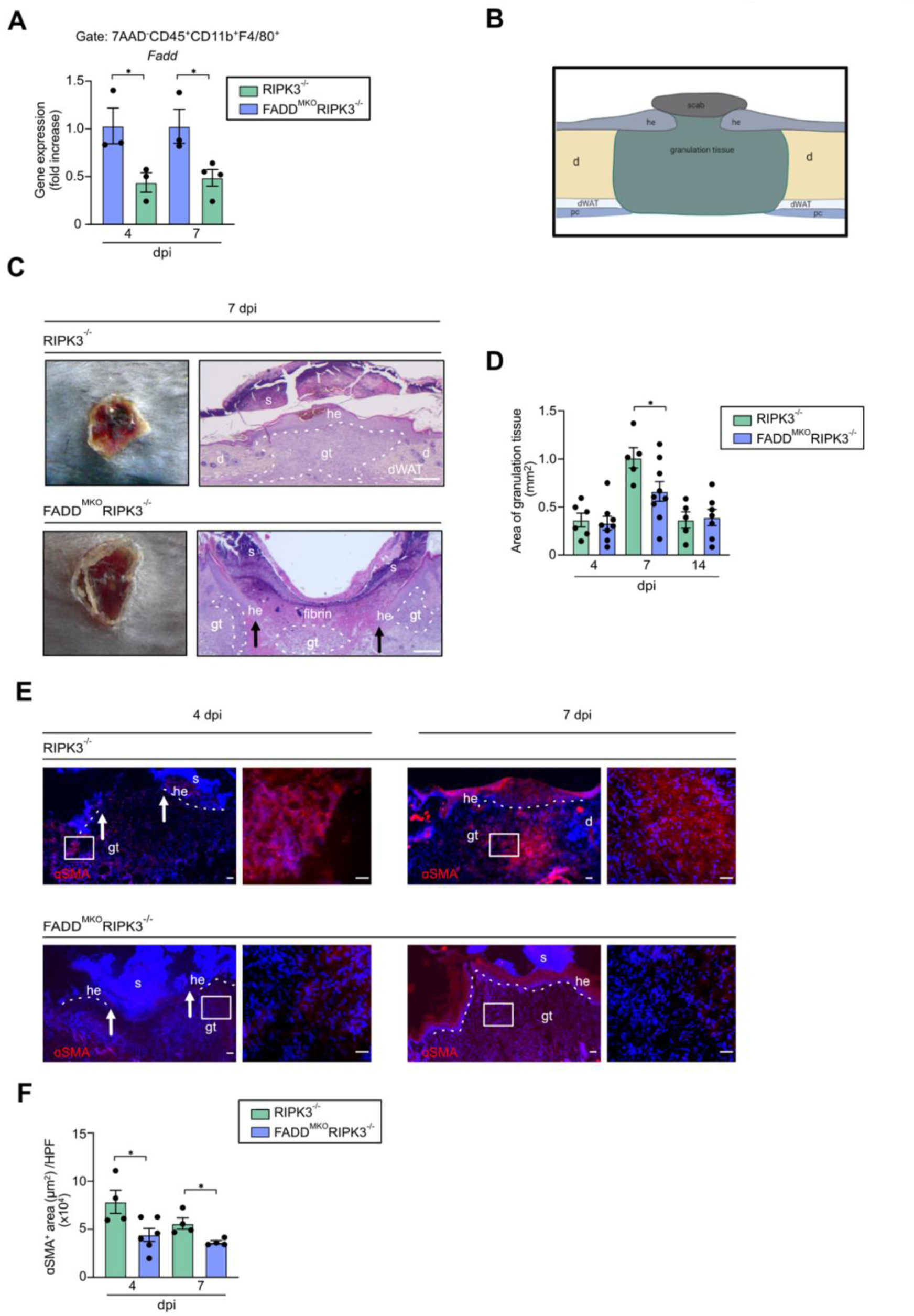
Modulating FADD-RIPK3 signaling in monocytes/macrophages impairs tissue repair. **A.** *Fadd* expression (fold-change) in RIPK3^-/-^ and FADD^MKO^RIPK3^-/-^ wound macrophages (7-AAD^-^CD45^+^CD11b^+^F4/80^+^) measured by qRT-PCR at 4 and 7 dpi. **B.** Scheme illustrating histology of excisional skin wound (he, hyper-proliferative epithelium; gt, granulation tissue; d, dermis; dWAT, dermal white adipose tissue; pc, panniculus carnosus; arrows indicate epithelial wound edge). **C.** Representative macroscopic (left) and H&E (right) images of wounds in mutant mice at 7 dpi; s, scab; scale, 200µm. **D.** Quantification of granulation tissue area (mm^2^) in mutant mice as indicated. **E.** Representative images of DAPI (blue) and ɑSMA (red) double immunostaining of wounds in mutant mice as indicated. Scale, 20µm. **F.** Quantification of myofibroblast differentiation (ɑSMA^+^ area/high power field (HPF)) in mutant mice as indicated. Mean value +/- SEM is represented. *p<0.05.

We then evaluated the impact of extrinsic apoptosis and necroptosis in the clearance of monocytes/macrophages during skin repair. We subjected FADD^MKO^RIPK3^-/-^ mice to excision skin injury hypothesizing that the absence of FADD and RIPK3 will prevent wound monocytes/macrophages from undergoing cell death. We analyzed wound healing dynamics and the capability to restore tissue architecture in FADD^MKO^RIPK3^-/-^ mice, measuring histomorphological features of repair and wound closure, such as epithelization, myofibroblast differentiation, and granulation tissue formation (**Fig. 2B**). Our findings revealed a significant reduction of granulation tissue formation in FADD^MKO^RIPK3^-/-^ mice compared to RIPK3^-/-^ mice at 7 dpi (**Fig. 2C, D**). In addition, quantitative immunofluorescence of ɑSMA staining, the signature marker for myofibroblast differentiation, revealed a significant reduction in FADD^MKO^RIPK3^-/-^ mice compared to the RIPK3^-/-^ mice at 4 and 7 dpi (**Fig. 2E, F**). These alterations are consistent with significant deficits during the early- and the mid-phase of repair, suggesting that FADD-RIPK3 signaling in wound monocytes/macrophages plays an important role in the early inflammatory and the granulation tissue-forming stage. Interestingly, when we compared wound healing dynamics in RIPK3^-/-^ mice with previously published data from our group in control mice, we found that modulating RIPK3 signalling alone did not lead to significant alterations (Coutelle, 2014; Willenborg, 2012; Willenborg 2021).

### Modulating FADD-RIPK3 signaling in macrophages leads to increased number and viability of Ly6C^high^ monocytes/macrophages

We investigated a functional relationship between delayed wound healing and extended viability of macrophages in FADD^MKO^RIPK3^-/-^ mice. Single-cell suspensions of wound tissues during subsequent phases of repair were analyzed by flow cytometry to quantify wound macrophages. No significant difference in relative and absolute number of wound macrophages in FADD^MKO^RIPK3^-/-^ and RIPK3^-/-^ mice was detected (**Fig. 3A, B**). In contrast, a significantly enriched pro-inflammatory Ly6C^high^ monocyte/macrophage population in FADD^MKO^RIPK3^-/-^ mice was identified when compared to the RIPK3^-/-^ mice over the entire time course of repair (**Fig. 3A, C**). The pro-inflammatory propensity of the wound environment in FADD^MKO^RIPK3^-/-^ mice was further shown by increased numbers of PMNs (7-AAD^-^CD45^+^CD11b^+^F4/80^-^Ly6G^+^) at 7 dpi when compared to RIPK3^-/-^ mice (**Fig. S3A, B**).

**Figure 3.**
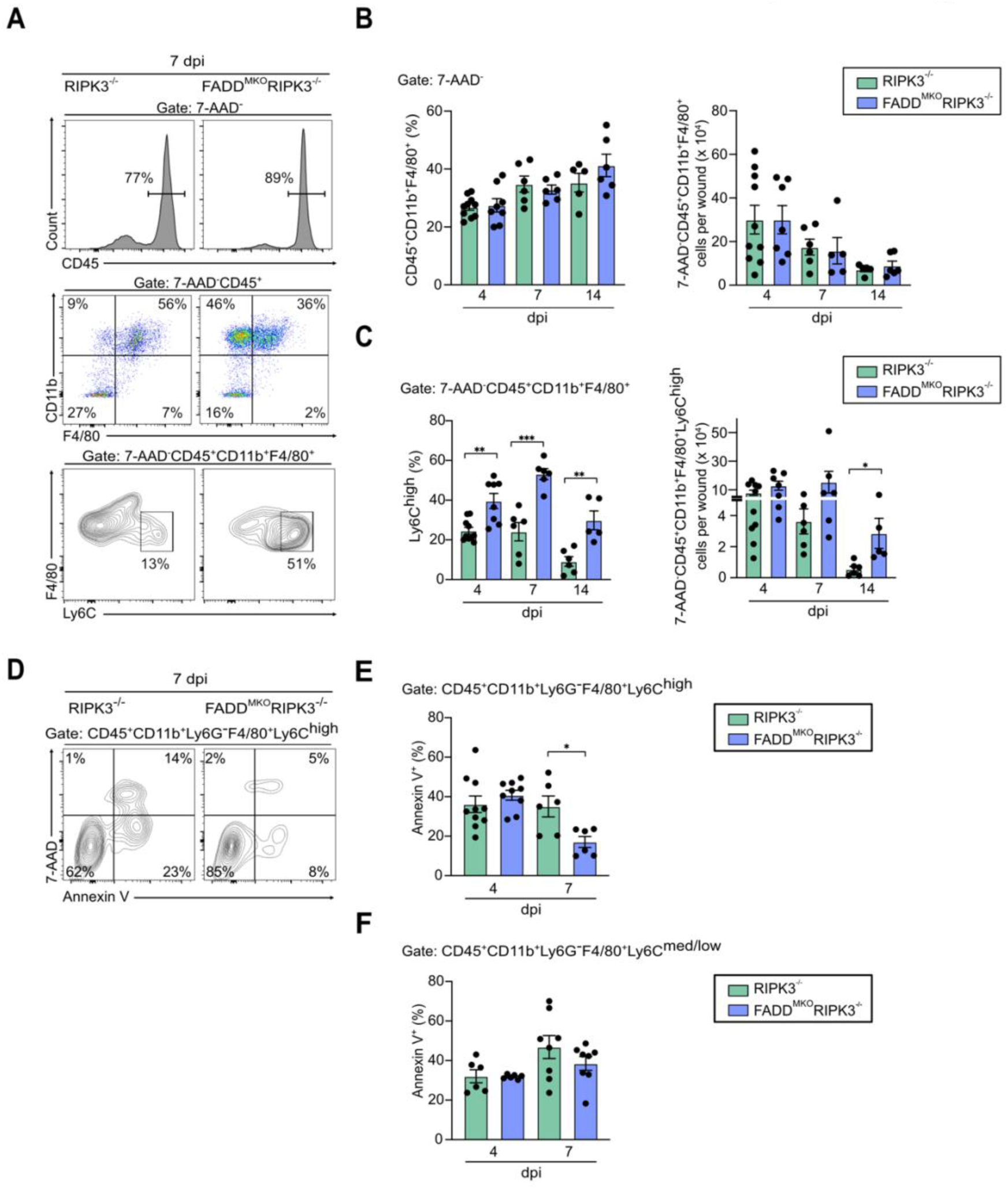
Modulating FADD-RIPK3 signaling in macrophages leads to an increased number and viability of Ly6C^high^ monocytes/macrophages. **A.** Representative 7 dpi FACS gating strategy used for identifying macrophages (7-AAD^-^ CD45^+^CD11b^+^F4/80^+^) and inflammatory macrophages (7-AAD^-^ CD45^+^CD11b^+^F4/80^+^Ly6C^high^) in wounds of mutant mice. **B.** Quantification of macrophage numbers in mutant mice as indicated. **C.** Quantification Ly6C^high^ macrophage numbers in mutant mice as indicated. **D.** Annexin V/7-AAD FACS profiles of Ly6C^high^ wound macrophages (7-AAD^-^CD45^+^CD11b^+^F4/80^+^Ly6C^high^) in mutant mice at 7 dpi. **E.** Quantification of Annexin V^+^Ly6C^high^ macrophages in mutant mice at 4 and 7 dpi. **F.** Quantification of Annexin V^+^Ly6C^med/low^ macrophages in mutant mice as indicated. Mean value +/- SEM is represented. *p<0.05; **p<0.01; ***p<0.001.

Next, we analyzed whether the Ly6C^high^ inflammatory monocyte/macrophage subpopulation showed an extended viability in wounds of FADD^MKO^RIPK3^-/-^ mice. To assess cell death, we subjected Ly6C^high^ wound monocytes/macrophages of FADD^MKO^RIPK3^-/-^ and RIPK3^-/-^ mice to 7-AAD and Annexin V double staining, followed by flow cytometric analysis. At 7 dpi, we observed a significant decrease of apoptotic (Annexin V^+^) Ly6C^high^ wound monocytes/macrophages in the FADD^MKO^RIPK3^-/-^ mice when compared to the RIPK3^-/-^ mice (**Fig. 3D, E**). Ly6C^medium/low^ wound monocytes/macrophages were not altered (**Fig. 3F**). These findings highlight an important role for the FADD-RIPK3 cell death pathway particularly in Ly6C^high^ monocyte/macrophages with critical consequences for tissue inflammation and resolution.

### Inhibiting TNF in FADD^MKO^RIPK3^-/-^ mice improves wound healing

To further characterize the activation state of wound macrophages in FADD^MKO^RIPK3^-/-^ mice, we quantified expression levels of established pro-inflammatory markers (*Tnfa*, *Il1b*, *Nos2*) in early wound macrophages using qRT-PCR. Our findings revealed a significantly higher expression of these genes in wound macrophages of FADD^MKO^RIPK3^-/-^ mice compared to macrophages of RIPK3^-/-^ and wild-type mice (**Fig. 4A**). Pro-inflammatory cytokines, in particular TNF, play a central role in driving inflammation and cell death in damaged tissues. Thus, we hypothesized a functional relationship between the sustained inflammation, excessive TNF and impaired healing dynamics in FADD^MKO^RIPK3^-/-^ mice.

**Figure 4.**
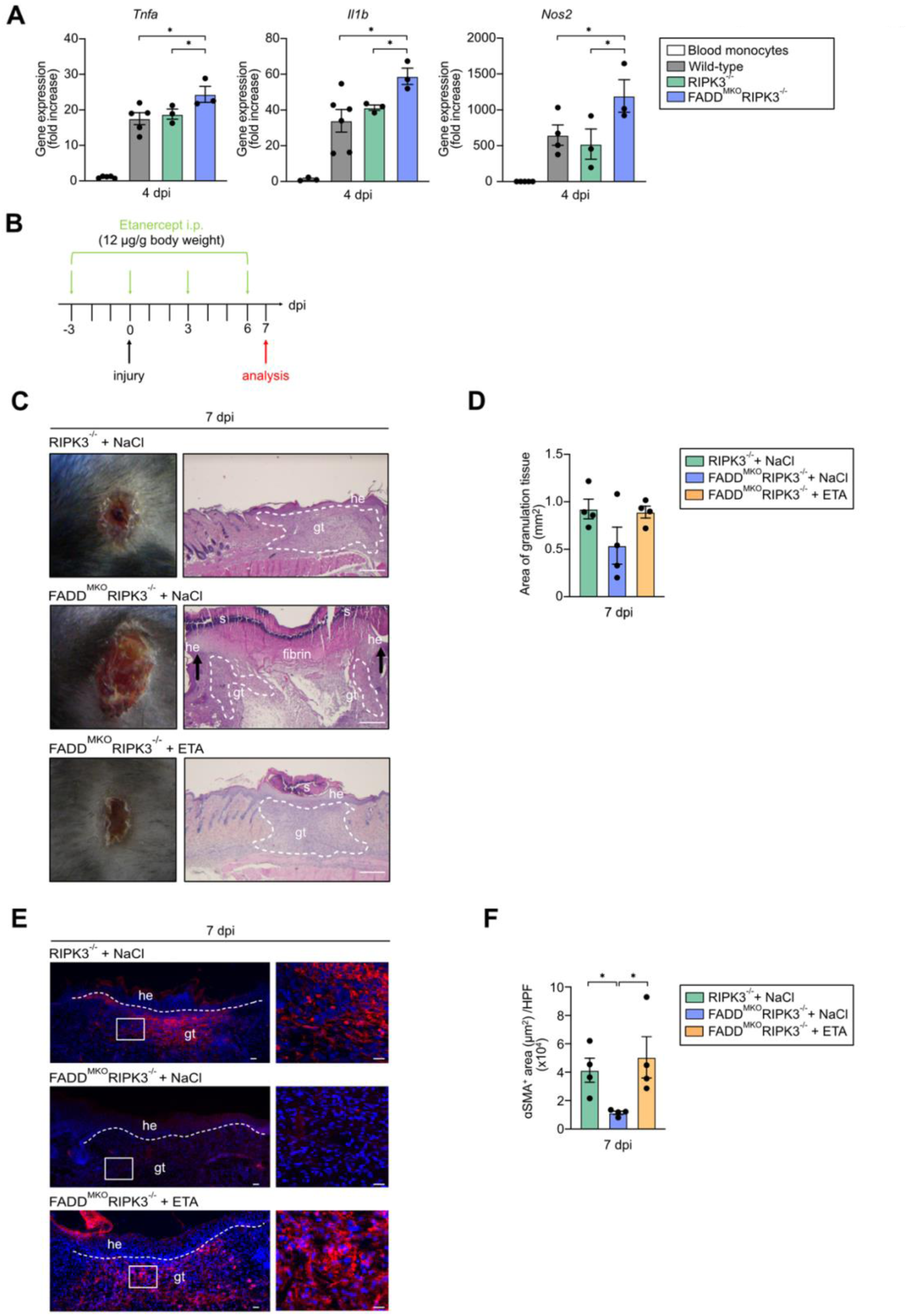
Inhibiting TNF in FADD^MKO^RIPK3^-/-^ mice improves wound healing. **A.** Gene expression in blood monocytes and in wound macrophages of wild-type, and mutant mice measured by qRT-PCR at 4 dpi. **B.** Etanercept (ETA) injection strategy. **C.** Macroscopic (left) and H&E (right) images of wounds of mutant mice receiving NaCl and FADD^MKO^RIPK3^-/-^ mice receiving ETA at 7 dpi. Scale, 200µm. **D.** Quantification of granulation tissue area (mm^2^) in mutant mice receiving NaCl or ETA at 7 dpi. **E.** Representative images of DAPI (blue) and ɑSMA (red) double immunostaining of wounds of mutant mice receiving NaCl or ETA at 7 dpi. Scale, 20µm. **F.** Quantification of myofibroblast differentiation (ɑSMA^+^ area/HFP) in mutant mice receiving NaCl or ETA at 7 dpi. he, hyper-proliferative epithelium; gt, granulation tissue. Mean value +/- SEM is represented. *p<0.05.

To investigate this hypothesis, we inhibited TNF in wounded FADD^MKO^RIPK3^-/-^ mice by using etanercept (ETA) (**Fig. 4B**). ETA is a soluble Fc:TNF receptor 2 (TNFR2) analog that binds and inhibits TNF, which is effective in the treatment of human TNF-mediated diseases such as psoriasis and rheumatoid arthritis (Weinblatt, 1999). Notably, the impaired wound healing response was restored by TNF inhibition in FADD^MKO^RIPK3^-/-^ mice. Quantitative analysis of H&E-stained wound sections and ɑSMA staining showed an increased granulation tissue formation in the FADD^MKO^RIPK3^-/-^ mice treated with ETA when compared to the FADD^MKO^RIPK3^-/-^ mice receiving NaCl (mock controls) (**Fig. 4C-F**). Taken together, these findings revealed that TNF-dependent signaling underlies the impaired healing response in FADD^MKO^RIPK3^-/-^ mice.

### TNF inhibition in FADD^MKO^RIPK3^-/-^ mice decreases the number of Ly6C^high^ wound monocytes/macrophages and activates pro-repair programs in late-stage wound macrophages

To elucidate the mechanism of how TNF attenuation in FADD^MKO^RIPK3^-/-^ mice improves wound healing, single-cell suspensions of wound tissues were analyzed by flow cytometry (**Fig. 5A**). Our findings revealed significantly reduced relative and absolute numbers of inflammatory Ly6C^high^ monocytes/macrophages in FADD^MKO^RIPK3^-/-^ mice receiving ETA when compared to mock controls (**Fig. 5B,C**). FADD deficiency in our model disables wound macrophages to undergo extrinsic apoptosis, therefore we speculated whether intrinsic apoptosis instead could contribute to the reduction of Ly6C^high^ wound monocytes/macrophages in FADD^MKO^RIPK3^-/-^ mice following ETA treatment. Interestingly, previously it was shown in an experimental rheumatoid arthritis model that anti-TNF treatment reduces the number of Ly6C^high^ monocytes/macrophages in inflamed joints through caspase-3- dependent apoptosis, while no significant increase in caspase-8 activation was observed (Huang, 2018). Here, we subjected wound single cell suspensions isolated from FADD^MKO^RIPK3^-/-^ mice treated with ETA and mock controls to 7-AAD and Annexin V double staining, followed by flow cytometric analysis. Notably, at 14 dpi, we observed significantly increased apoptosis (Annexin V^+^) in Ly6C^high^ monocytes/macrophages in FADD^MKO^RIPK3^-/-^ mice treated with ETA when compared to mock controls (**Fig. 5D,E**). Unexpectedly, at 7 dpi we did not observe a similar trend, suggesting that different mechanisms than apoptosis contribute to the reduction of inflammatory monocytes/macrophages at the mid-stage of repair (**Fig. 5C-E**).

**Figure 5.**
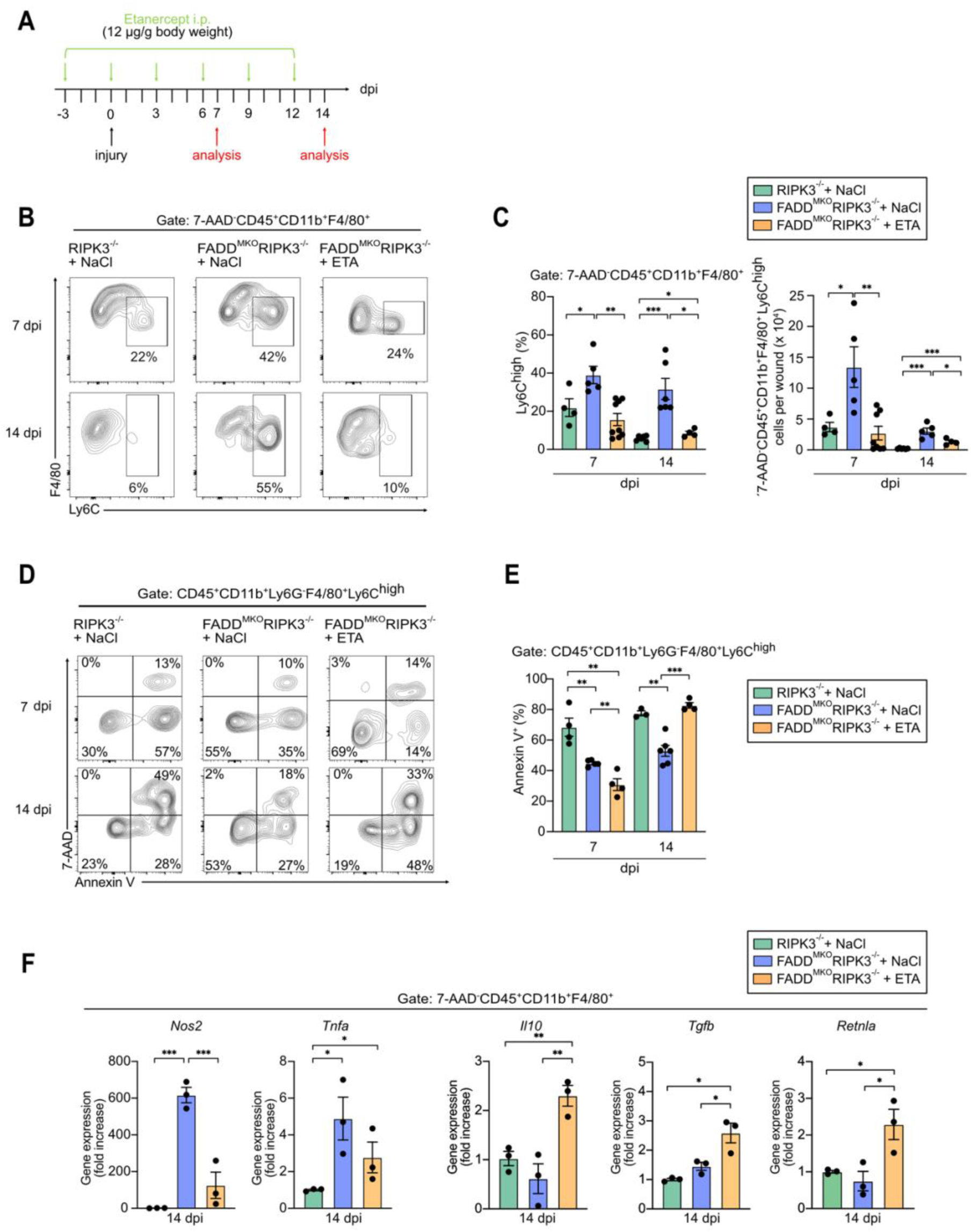
TNF inhibition in FADD^MKO^RIPK3^-/-^ mice decreases the number of Ly6C^high^ wound monocytes/macrophages and increases pro-repair programs in late-stage wound macrophages. **A.** ETA injection strategy. **B.** Representative 7 and 14 dpi FACS gating strategy for inflammatory macrophages (7-AAD^-^ CD45^+^CD11b^+^F4/80^+^Ly6C^high^) in wounds of mutant mice treated with NaCl or ETA. **C.** Quantification of Ly6C^high^ macrophages at 7 and 14 dpi in wounds of mutant mice treated with NaCl or ETA. **D.** Annexin V/7-AAD FACS profiles of Ly6C^high^ wound macrophages (7-AAD^-^CD45^+^CD11b^+^F4/80^+^Ly6C^high^) in wounds of mutant mice treated with NaCl or ETA at 7 and 14 dpi. **E.** Quantification of Annexin V^+^Ly6C^high^ macrophages in the wounds of mutant mice treated with NaCl or ETA. **F.** Gene expression in sorted wound macrophages (7-AAD^-^CD45^+^CD11b^+^F4/80^+^) of mutant mice treated with NaCl or ETA measured by qRT-PCR. Mean value +/- SEM is represented. *p<0.05, **p<0.01, ***p<0.001.

We then addressed the question whether repair programs in late-stage wound macrophages are perturbed in FADD^MKO^RIPK3^-/-^ mock controls and restored upon TNF blockade. We sorted wound macrophages of RIPK3^-/-^ and FADD^MKO^RIPK3^-/-^ mice treated with NaCl, and FADD^MKO^RIPK3^-/-^ mice treated with ETA at 14 dpi and analyzed the expression of established pro-inflammatory and pro-repair genes using qRT-PCR. Our analysis revealed that wound macrophages of RIPK3^-/-^ mice behaved similarly when compared to wild-type mice i.e., they showed down-regulated pro-inflammatory genes (*Nos2* and *Tnfa*) and up-regulated pro-repair genes (*Il10, Tgfb, Retnla*) in the late-stage, consistent with previous observations (Knipper, 2015; Willenborg, 2021). In contrast, wound macrophages of FADD^MKO^RIPK3^-/-^ mock controls, showed a reversed polarization state, with significantly higher expression of pro-inflammatory genes and a tendency of reduced pro-repair genes when compared to RIPK3^-/-^ mice (**Fig. 5F**). Notably, anti-TNF treatment reverted the perturbed wound macrophage gene expression dynamics in FADD^MKO^RIPK3^-/-^ mice, by a down-regulation of pro-inflammatory genes and an up-regulation of pro-repair genes when compared to both mock controls and RIPK3^-/-^ mice. Interestingly, flow cytometric analysis of wound macrophages at 14 dpi revealed increased relative and absolute numbers in FADD^MKO^RIPK3^-/-^ mice receiving ETA when compared to RIPK3^-/-^ mice (**Fig. S4A, B**). Together, these findings suggest a multitasking role for TNF in wound monocyte/macrophage function in healthy conditions: beyond regulating cell survival, TNF controls cell death and surveilles timely activation of late-stage repair programs. Our findings suggest that TNF blockade in FADD^MKO^RIPK3^-/-^ mice improves wound healing through a combination of mechanisms.

## Discussion

Resolution of inflammation is of fundamental importance to ensure healing and restoration of tissue homeostasis after injury. Different mechanisms contribute to resolution of inflammation in damaged and inflamed tissues (Chen, 2015; Knipper, 2015; Bosurgi, 2017; Zhang, 2019). The current view is that in most repairing tissues, including the skin, inflammatory monocytes/macrophages (Ly6C^high^) undergo a phenotypic switch from a pro-inflammatory towards an anti-inflammatory phenotype (Willenborg, 2012; Knipper, 2015; Willenborg, 2021; Sanin, 2022). In addition, monocytic cells can be cleared through lymphatic drainage (Harmsen, 1985; Bellingan, 1996), recently described trans-differentiation (Meng, 2016; Sinha, 2018; Haider, 2019; Li, 2021), and/or cell death (Desmoulière, 1995; Kuhlmann, 2001; Janssen, 2011; Gautier, 2013; Moriwaki, 2014; Wu, 2014; Bleriot, 2015; Legrand, 2019). However, the contribution and the dynamics of these different resolution mechanisms, particularly also the role of different cell death modalities in the removal of inflammatory macrophages in inflamed tissues remains elusive. This is an important and timely question, in light of emerging findings demonstrating the importance of distinct modes of cell death in regulating tissue inflammation and resolution (Gautier, 2013; Pasparakis, 2015; Pang, 2021).

Here, we demonstrate in a model of mechanical skin injury that FADD-caspase-8- mediated apoptosis plays an important role in the removal of wound macrophages across all stages of physiological repair. Our findings suggest that extrinsic apoptosis mediated by FADD plays in particular a critical role in the timely removal of a potent Ly6C^high^ pro-inflammatory subset of macrophages. Failure to clear this pro-inflammatory macrophage subpopulation in FADD^MKO^RIPK3^-/-^ mice, leads to monocyte/macrophage persistence, which is associated with impaired wound healing characterized by prolonged inflammation, decreased myofibroblast differentiation and reduced granulation tissue formation. Our findings also suggest that FADD-RIPK3- deficient Ly6C^high^ wound monocytes/macrophages are unable to employ alternative mechanisms for timely clearance. We observed no major effects in wound healing dynamics in *Ripk3*^-/-^ mice, suggesting that necroptosis itself does not play a critical role in the removal of monocytes/macrophages in physiological wound healing under specific pathogen free conditions in healthy mice.

We show that relative and absolute numbers of Ly6C^high^ monocytes/macrophages are increased in wounds of FADD^MKO^RIPK3^-/-^ mice. We hypothesize that the delayed clearance and prolonged persistence of this pro-inflammatory macrophage population contributes to the delayed wound healing in FADD^MKO^RIPK3^-/-^ mice. Our findings suggest that the prolonged persistence of Ly6C^high^ macrophages promotes a highly inflammatory wound environment, characterized by increased expression of pro-inflammatory mediators and increased PMN recruitment, both of which counteract the emergence of pro-repair macrophages.

We showed that persisting inflammatory macrophages in wounds in FADD^MKO^RIPK3^-/-^ mice were characterized by increased TNF expression. Beside its pro-inflammatory activities, TNF has a central role in promoting the survival of inflammatory macrophages (Lombardo, 2007). We propose that Ly6C^high^ monocyte/macrophage viability is prolonged when TNF-triggered cell death pathways are inhibited in wounded FADD^MKO^RIPK3^-/-^ mice. Thereby, having increased numbers of wound macrophages that produce more TNF, likely the wound environment becomes more inflammatory. This cascade then leads to a vicious circle in which TNF fuels the growth/differentiation of inflammatory wound macrophages that express more TNF. Thus, indicating that unbalanced TNF activities are one of the key mediators driving the enhanced inflammatory wound environment in mutant mice.

Besides TNF, we also observed an increased expression of *Nos2* in wound macrophages of FADD^MKO^RIPK3^-/-^ mice, both in early and late stage of wound healing. Macrophage iNOS is a major source of nitric oxide (NO) production in inflamed tissues. NO is known to repress oxidative phosphorylation in macrophages, thus inhibiting “M2”-like pro-repair polarization both *in vitro* and *in vivo* (Baseler, 2016; Palmieri, 2020). Moreover, NO production in macrophages has been shown to sensitize macrophages to cell death during bacterial infection (Mariott, 2004; Zwaferink, 2008), speculating that NO-producing macrophages are primarily cleared through cell death. Together, these findings suggest that impaired wound healing in FADD^MKO^RIPK3^-/-^ mice is caused by the prolonged TNF-driven viability of Ly6C^high^ monocytes/macrophages and the combined suppressive effect of excessive NO and TNF on switching inflammatory macrophages into pro-repair mode.

TNF inhibition improved wound healing in FADD^MKO^RIPK3^-/-^ mice. These findings highlight a functional relationship between excessive TNF and the maladaptive tissue repair response in FADD^MKO^RIPK3^-/-^ mice. Our findings propose different mechanisms how TNF blockade might rescue impaired healing in FADD^MKO^RIPK3^-/-^ mice. First, following ETA administration, we observed a significant decrease of absolute Ly6C^high^ monocyte/macrophage numbers and increased Ly6C^low/medium^ macrophages in the mid- and late stage of repair. Given the pro-survival mechanism of TNF signaling, we suggest that TNF inhibition could increase the onset of intrinsic apoptosis of Ly6C^high^ monocytes/macrophages, as shown recently in an experimental model of rheumatoid arthritis (Huang, 2018). Thereby, the vicious circle of uncontrolled TNF release from inflammatory macrophages and its contribution to an inflamed and hostile wound environment is broken by etanercept. Secondly, we demonstrate that TNF blockade in FADD^MKO^RIPK3^-/-^ mice is associated with attenuated expression of pro-inflammatory mediators and enhanced expression of pro-repair programs in late-stage wound macrophages. Recently, we reported similar effects of TNF blockade in wound healing in healthy mice, indicating that TNF suppressive-M2 effects are also present in FADD^MKO^RIPK3^-/-^ mice (Dichtl, 2022). Third, we propose that improved wound healing in FADD^MKO^RIPK3^-/-^ mice after ETA treatment is achieved not only through the reactivation of the suppressed repair phenotype, but also through an increased absolute number of late-stage pro-repair wound macrophages. Together, these findings unravel the complexity of TNF activities in tissue repair and suggest a combined effect of different mechanisms of how TNF blockade improves wound healing in FADD^MKO^RIPK3^-/-^ mice.

Finally, anti-TNF therapies are a core anti-inflammatory approach for chronic inflammatory skin diseases that resemble unsuccessful wound healing responses including psoriasis (Köbner phenomenon), ulcerative lesions caused by pyoderma gangraenosum or necrobiosis lipoidica. However, the mechanistic understanding of the repair-promoting effects of anti-TNF therapies in these pathological conditions is not entirely resolved. Our findings contribute to a better mechanistic understanding of repair-promoting effects of anti-TNF therapies and ultimately guide the development of therapeutic approaches to ameliorate wound healing disorders in patients.

## Acknowledgements

We thank Michael Piekarek and Gabriele Scherr for excellent technical assistance; Dr. Gunter Rappl (Central Cell Sorting Facility, CMMC, Cologne University) for specialized technical support; project leaders and central projects of the CRC 1403 for helpful discussions. This work was supported by the Deutsche Forschungsgemeinschaft (DFG, German Research Foundation): the CRC 1403 (project number 414786233 to SAE, MP, HK) and the Germany’s Excellence Strategy – CECAD (EXC 2030 – 390661388 to SAE, MP, HK); the Center for Molecular Medicine (to SAE, MP, HK).

## Author contributions

LI, SW and SAE wrote the manuscript; LI und SW designed and performed most of the experiments and analyzed the data; DW and SW performed and analyzed data in wild-type mice; MP and HK provided mutant mouse models, planned experiments, discussed data and critically revised the manuscript; SAE conceived and supervised the study; all authors commented on and edited the manuscript.

