## Supplementary Material for "FADD- and RIPK3-mediated cell death ensures timely clearance of wound macrophages and promotes wound healing"

**Figure S1 related to Fig. 1.** Gating strategy used for identifying macrophages ($CD45^+CD11b^+Ly6G^-F4/80^+$) and inflammatory macrophages ($CD45^+CD11b^+Ly6G^-F4/80^+Ly6C^{high}$) in 7-AAD and Annexin V co-staining assay.

**Figure S2 related to Fig. 2. A.** Schematic illustration of *Fadd* deletion and genotyping strategy (tail biopsies); E, exon; del, deletion; fl, flox; bp, base pair. **B.** Mendelian distribution. **C.** *Ripk3* expression in wild-type, $RIPK3^{-/-}$ and $FADD^{MKO}RIPK3^{-/-}$ wound macrophages ($7\text{-}AAD^-CD45^+CD11b^+F4/80^+$) measured by qRT-PCR at 4 and 14 dpi. **D.** Gating strategy used to identify lymphocytes ($7\text{-}AAD^-CD45^+CD11b^-SSC^{low}$), PMNs ($7\text{-}AAD^-CD45^+CD11b^+SSC^{high}Ly6G^+$) and $Ly6C^{high}$ monocytes ($7\text{-}AAD^-CD45^+CD11b^+SSC^{low}Ly6C^{high}$) in peripheral blood depleted from red blood cells. **E.** Distribution of lymphocytes, PMNs and $Ly6C^{high}$ monocytes in the blood of $RIPK3^{-/-}$ and $FADD^{MKO}RIPK3^{-/-}$ mice. Mean value +/- SEM is represented.

**Figure S3 related to Fig. 3. A.** Ly6G FACS profiles of $RIPK3^{-/-}$ and $FADD^{MKO}RIPK3^{-/-}$ $7\text{-}AAD^-CD45^+CD11b^+F4/80^-$ wound cells isolated at 4, 7 and 14 dpi. **B.** Quantification of $7\text{-}AAD^-CD45^+CD11b^+F4/80^-Ly6G^+$ wound cells (PMNs) in $RIPK3^{-/-}$ and $FADD^{MKO}RIPK3^{-/-}$ mice at 4, 7 and 14 dpi. Mean value +/- SEM is represented. *$p<0.05$.

**Figure S4 related to Fig. 5. A.** CD11b and F4/80 FACS profiles of wounds of mutant mice treated with NaCl or ETA at 7 and 14 dpi. **B.** Quantification of macrophages ($7\text{-}AAD^-CD45^+CD11b^+F4/80^+$) in the wounds of mutant mice treated with NaCl or ETA at 7 and 14 dpi. Mean value +/- SEM is represented. *$p<0.05$, **$p<0.01$, ***$p<0.001$.

**Injarabian *et al.* 2022 Figure S1**

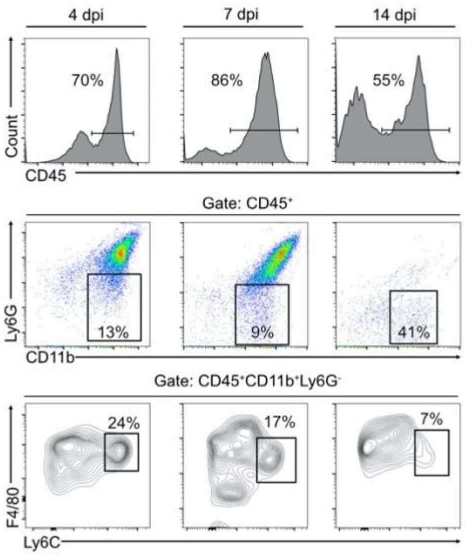

**Injarabian *et al.* 2022 Figure S2**

**A**

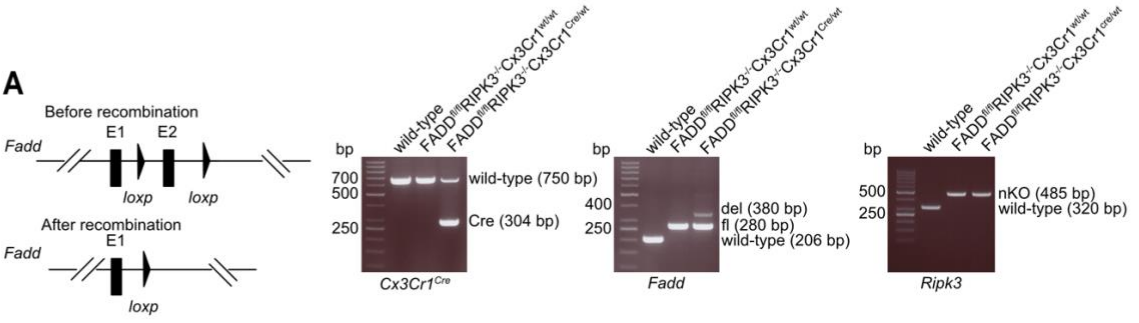


**B**

| Genotype | Number of mice (%) | % Expected |
|---|---|---|
| $FADD^{fl/fl}RIPK3^{-/-}Cx3Cr1^{wt/wt}$ | 55 (47) | 50 |
| $FADD^{fl/fl}RIPK3^{-/-}Cx3Cr1^{Cre/wt}$ | 61 (53) | 50 |

**C**

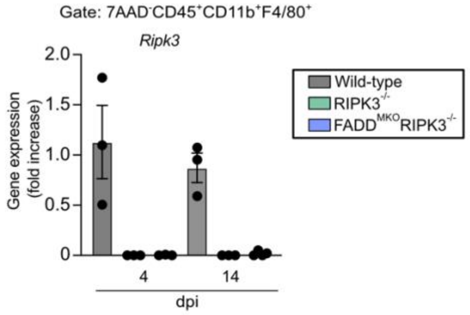


**D** **E**

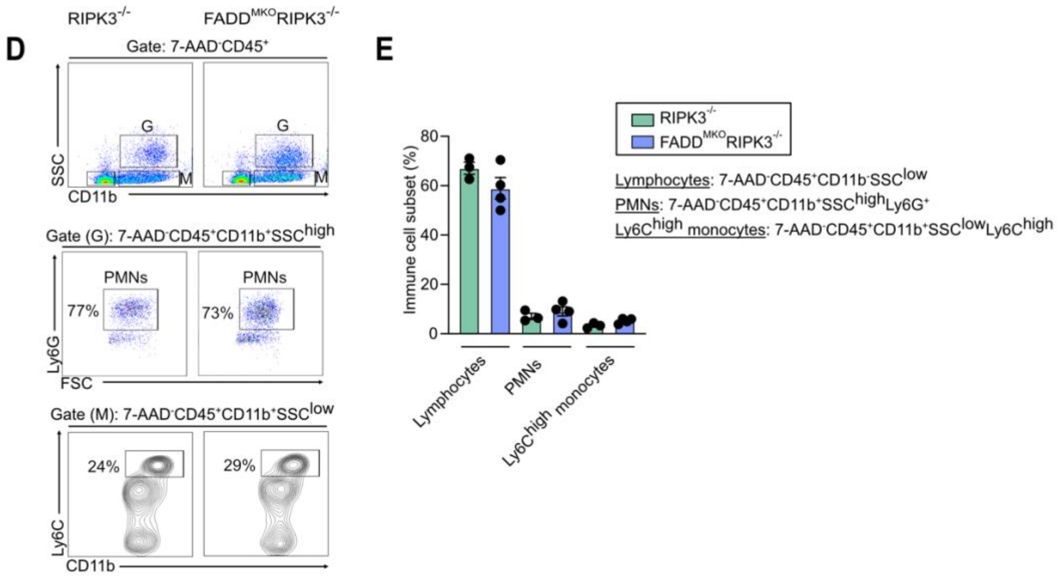

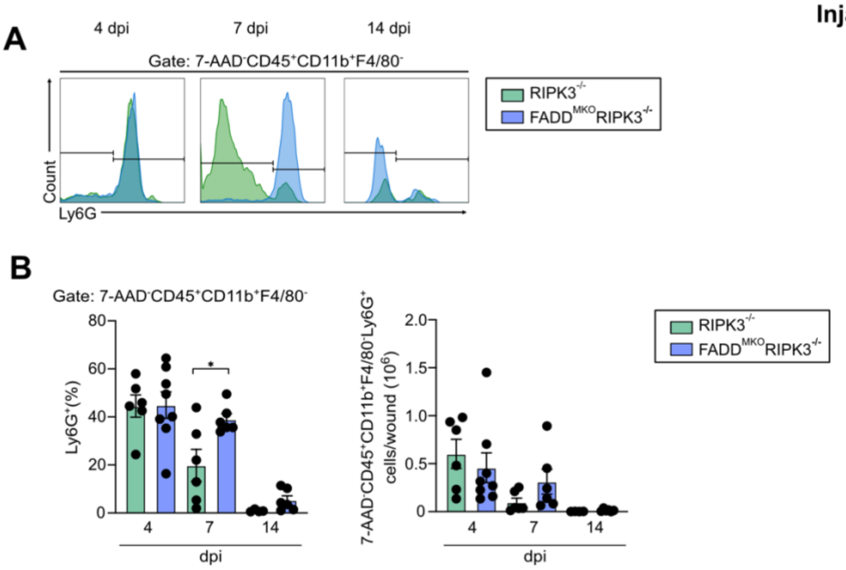
A
4 dpi
7 dpi
14 dpi
Gate: 7-AAD-CD45+CD11b+F4/80-
RIPK3-/-
FADDMKORIPK3-/-
Count
Ly6G
B
Gate: 7-AAD-CD45+CD11b+F4/80-
Ly6G+(%)
80
60
40
20
0
4
7
14
dpi
*
7-AAD-CD45+CD11b+F4/80-Ly6G+ cells/wound (106)
2.0
1.5
1.0
0.5
0
RIPK3-/-
FADDMKORIPK3-/-

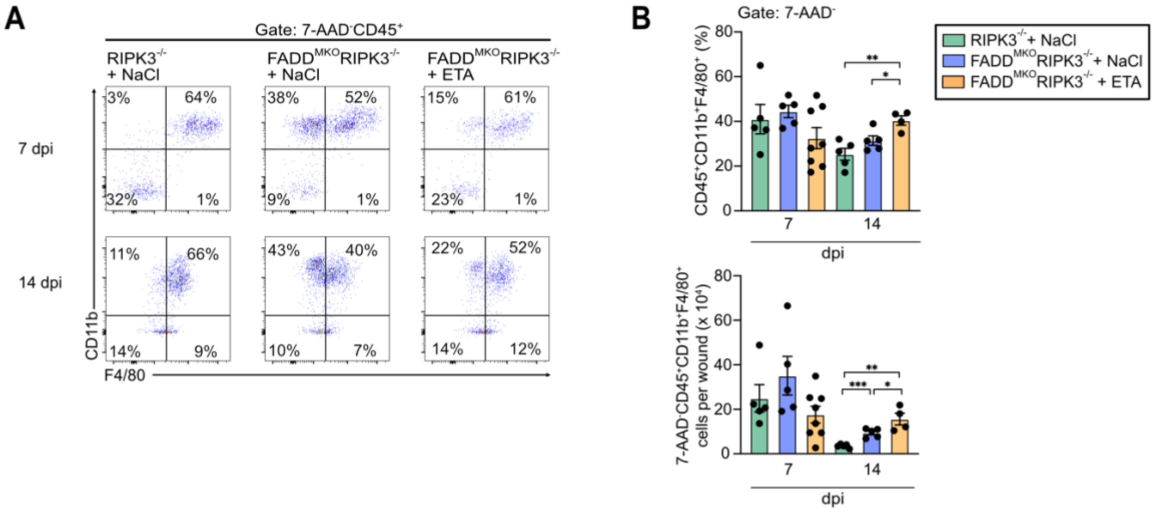
A
Gate: 7-AAD-CD45+
RIPK3-/- + NaCl
FADDMKORIPK3-/- + NaCl
FADDMKORIPK3-/- + ETA
7 dpi
14 dpi
3%
64%
32%
1%
38%
52%
9%
1%
15%
61%
23%
1%
11%
66%
14%
9%
43%
40%
10%
7%
22%
52%
14%
12%
CD11b
F4/80
B
Gate: 7-AAD-
CD45+CD11b+F4/80+ (%)
7-AAD-CD45+CD11b+F4/80+ cells per wound (x 10⁴)
dpi
RIPK3-/- + NaCl
FADDMKORIPK3-/- + NaCl
FADDMKORIPK3-/- + ETA
